# The ratio between experimental and theoretical localization precision of single molecules is sensor-dependent and typically larger than one

**DOI:** 10.1101/2024.10.10.617567

**Authors:** Alfonso Brenlla-Lopez, Laila Deen, Paolo Annibale

**Affiliations:** School of Physics and Astronomy, University of St Andrews, United Kingdom

## Abstract

Since the advent of stochastic localization microscopy approaches in 2006, the number of studies employing this strategy to investigate the sub-diffraction limit features of fluorescently labeled structures in biology, biophysics and solid state samples has increased exponentially. Underpinning all these approaches is the notion that the position of single molecules can be determined to high precision, provided enough photons are collected. The determination of exactly how precisely, has been demanded to formulas that try to approximate the so-called Cramer Rao Lower Bound based on input parameters such as the number of photons collected from the molecules, or the size of the camera pixel. These estimates should however be matched to the experimental localization precision, which can be easily determined if instead of looking at single beads, we study the distance between a pair. We revisit here a few key works, observing how these theoretical determinations tend to routinely underestimate the experimental localization precision, of the order of a factor two. A software-independent metric to determine, based on each individual setup, the appropriate value to set on the localization error of individual emitters is provided.

## Introduction

The idea that a single molecule emitter can be localised with an accuracy much higher than the diffraction limit is now a widespread notion and the cornerstone of fluorescence localization microscopy methods. This has spawned a large number of studies investigating cellular structures at high resolution(Lelek et al. 2021) and it is now proposed that these approaches can begin to complement ultrastructural approaches, e.g. electron microscopy, for structural biology(Liu, Lavis, and Betzig 2015).

Formulas that calculate the localization accuracy of single emitters imaged under a microscope and onto a camera sensor are based on the optical setup parameters and the number of photons collected. An original formulation was first proposed in 2002 by Thompson et al(Thompson, Larson, and Webb 2002), which developed an earlier theory by Bobroff(Bobroff 1986). Thompson’s work proposed a formula to estimate the localization precision of a point emitter, and an associated computational algorithm for the fitting, based on the use of Gaussian masks. Thompson formula contains effectively three terms: one to account for photon noise, one to correct for the finite size of the pixel, and the third term to take into account background noise for the case of weak emitters. Photon noise scales as 1/N with respect to the total number of collected photons, whereas background noise scales as 1/N^2^. This resulted in the well-known equation:

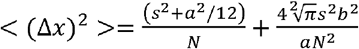

where *a* defines the pixel size at the sample, and *b* the background photons measured in the fitting window of the given emitter. This formula was incorporated in the original manuscripts that in 2006 proposed PALM(Betzig et al. 2006), fPALM(Hess, Girirajan, and Mason 2006) and STORM(Rust, Bates, and Zhuang 2006). Simply put, the formula shows clearly that the more photons are collected, the lower the localization error is. The formula was then used widely throughout the first years of localization microscopy research; nonetheless, a few years later, Mortensen et al (Mortensen et al. 2010) suggested to revisit the formula, implying that Thompon’s formula overestimated the localization error. Mortensen approach introduced a more precise localization scheme and evaluated Thompson’s error estimate as 1.78 times larger when fitting single emitters with Gaussian Masks (Figure 1) over Maximum Likelihood Estimation using Gaussians (MLEwG), suggesting that MLEwG localization would allow getting closer to the theoretical lower localization error, known as the Cramer-Rao Lower Bound (CRLB) (Small and Stahlheber 2014). Based on this limit, in the presence of an adequate number of photons, sub-nanometer localizations can be achieved on bright, isolated emitters, in principle allowing to study molecular interactions on spatial scales previously accessible only to single-molecule FRET.

**Figure 1.**
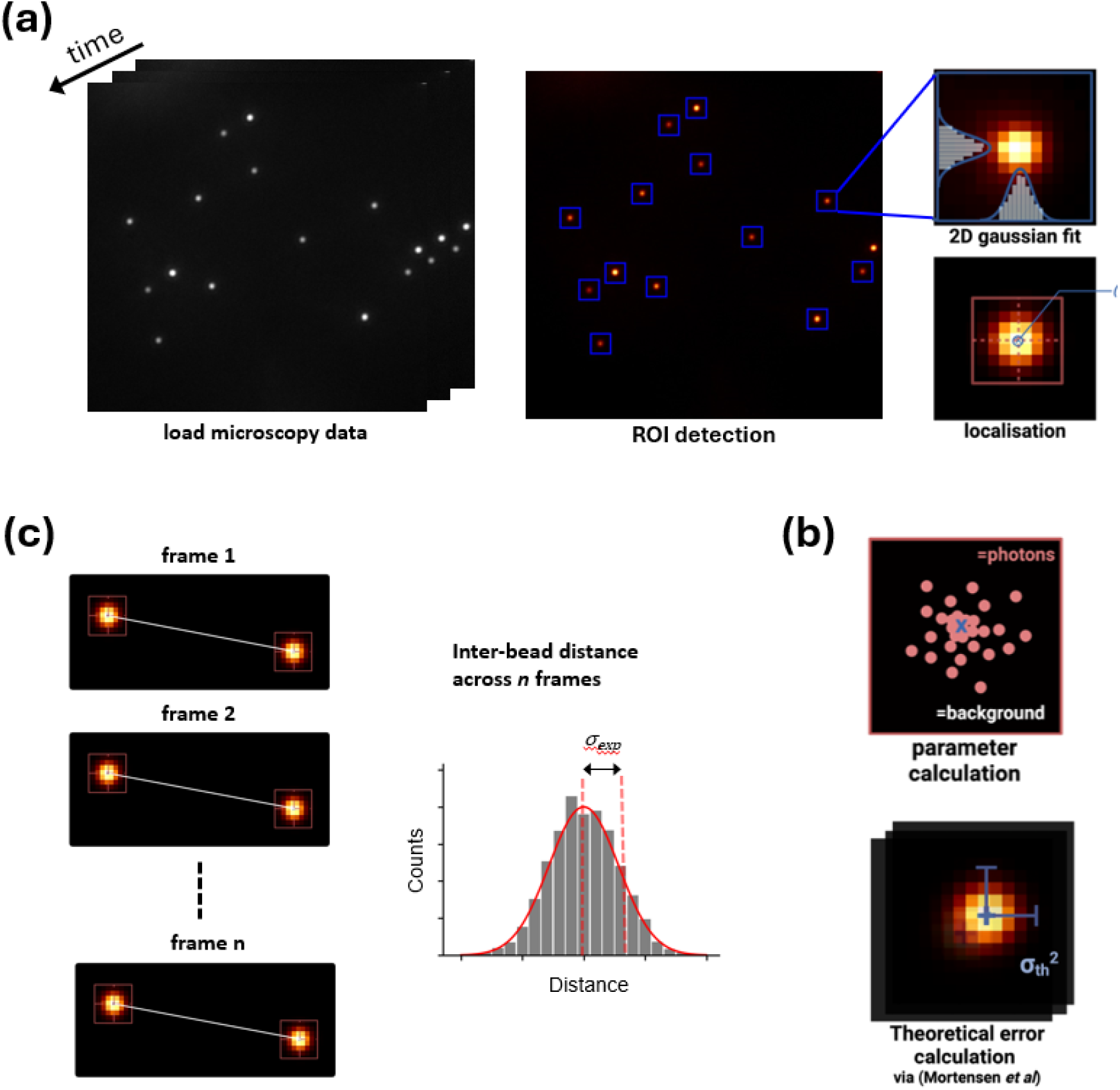
Experimental localization error from pairwise distances of single bright fluorescent emitters. **(a)** Regions of interest (ROI) with signal above a certain threshold are identified and fitted with a 2D Gaussian PSF to yield centroid coordinates x_m_ and y_m_. **(b)** The fit parameters from **(a)** also include the number of photons in the image, which are used to calculate the theoretical localisation error. **(c)** The sub-pixel centroid coordinates x_m_ and y_m_ enable the determination of interbead distances over all frames and the experimental localisation error.

In practice, however, colocalizations in the nm range have proved elusive, owing to the intrinsic nanoscale fluctuations of any microscopy setup as well as the pixelated nature of the detector and the noise characteristic of the camera electronics (Pertsinidis, Zhang, and Chu 2010). In this respect, it is important to note that Mortensen et al., who measured the fluctuations in the distance between pairs of 40 nm immobilized beads(Mortensen et al. 2010), observed that the variance of the measured (experimental) distances appeared to be always twice the one expected from the CRLB, which was attributed to the effect Electron Multiplication gain of the EMCCD camera (the authors used an Andor iXon DV887 back-illuminated camera, relying on a e2V CCD97 chip). At the same time, forward scattering interferometry measurements of immobilized beads on glass surfaces allowed measuring their distance with a precision better than 0.5 nm, indicating that no fundamental problem exists in achieving the beads and setup stability necessary for such a precision(Nugent-Glandorf and Perkins 2004).

This raises the question as to whether the localization precision estimate broadly used in the field, and which is used to render most single molecule-based super-resolution microscopy images, is an accurate estimator of the actual real-life localization precision. We shall note here that methods have been proposed to generate an estimate of the localization precision in superresolution datasets from the datasets themselves (Descloux, Grussmayer, and Radenovic 2019), (Endesfelder et al. 2014), albeit these approaches would incorporate any variability from sample preparation (e.g. cell chemical fixation), and therefore do not necessarily convey the fundamental resolution achievable with a given setup.

## Results and discussion

In order to address this question, we used an EMCCD camera having a comparable sensor (Photometrics Cascade II 512) to the one used in the original Mortensen manuscript and acquired images of sub-diffraction limited diameter (100 nm) fluorescent beads (**Figure 1a**). We reasoned that a direct measurement of the beads localization precision can be obtained from the standard deviation of the repeated measured distance between any two beads. Based on the notion, due to error propagation, that the standard deviation of the measured distance between any pair of beads arises from the sum in quadrature of the localization precision of each individual bead 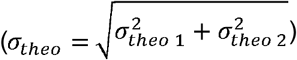, we compared the experimental localization precision of a pair of beads (σ_exp_) (**Figure 1b**) to the sum in quadrature of the beads localization precision calculated according to Mortensen et al (σ_theo_) (**Figure 1c**). Ideally, if the localization precision formula reported in the literature is accurate, the two values should match.

To fit the beads position (x_m_, y_m_) we used the MLEwG algorithm and code reported by the authors(Mortensen et al. 2010) (**Figure 2a-b**), as well as a Least Squares approach, which allowed us measuring bead-to-bead distances over movies containing one thousand frames. The Gaussian fits to the data proved to be -at least qualitatively-satisfactory, as shown by the lack of obvious features in the residual plots (**Figure S1**). The inter-bead distance **(Figure 2c)** oscillates around a central value over time and follows a Gaussian distribution whose standard deviation corresponds to the experimental localization error (σ_exp_) for that given inter-bead distance. Both experimental and theoretical localization errors over all bead pairs follow Gaussian distributions **(Figure 2d)**, with both variance and mean values larger for the experimental set.

**Figure 2.**
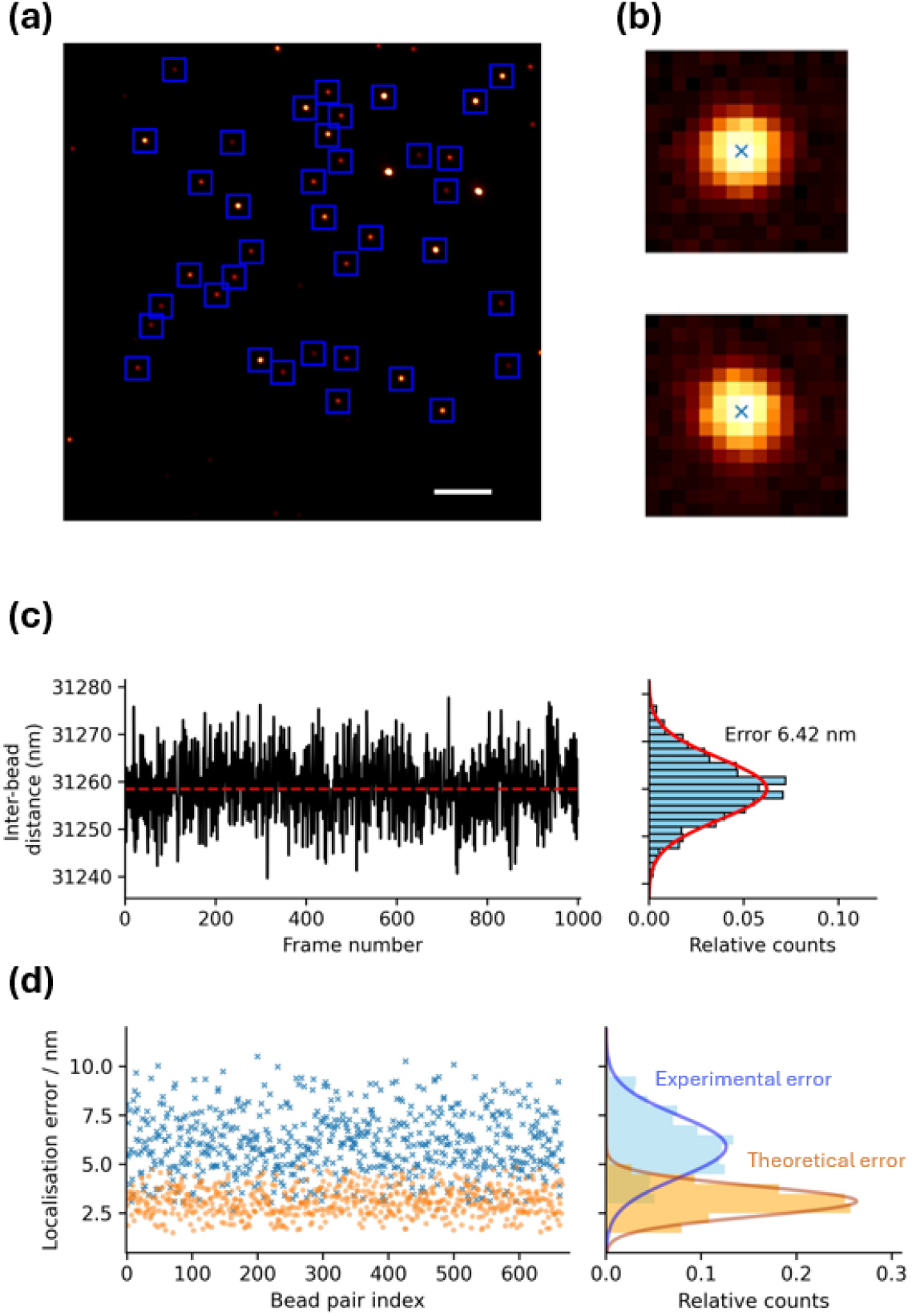
Experimental and theoretical errors for an EMCCD Cascade camera with gain disabled. **(a)** Field of view with regions of interest containing individual beads marked as blue squares **(b)** Centroid positions (blue cross) obtained via MLE fitting for two different beads. **(c)**. Distance over time for the two beads shown in (b) and the corresponding histogram with a Gaussian fit shown in red. **(d)**. Experimental and theoretical localisation errors for all the bead pairs show in (a) and their corresponding histograms

The ratios between experimental (σ_exp_) and theoretical localization (σ_theo_) for the bead pairs are therefore not the same, and it appears that σ_theo_ underestimates significantly the actual localization error of the individual beads, albeit the relationship between the two is linear (**Figure S2**). In order to understand better the source of this discrepancy, we explored how this metric changed when using different settings or sensors. The first observation is that the errors ratio varies depending on the sensor used **(Figure 3)**. The Cascade camera has two readout amplifiers that can be swirchen by software, one with electron charge multiplication gain (emGain) and another for traditional readout without emGain. The presence of these two readout modes within the same sensor allows for a direct measurement of any effect of emGain on the data noise and thus the localisation error. With emGain enabled, the ratio between experimental and theoretical error for the Cascade camera is ∼2.4 (**Figure 3b**), while disabling emGain brings the ratio down to ∼1.9 (**Figure 3a**). Notably, this appears to suggests that the factor of 2 in the ratio σ_exp_ / σ_theo_ reported by Mortensen does not arise from additional noise inherent to the electron multiplication process.

**Figure 3.**
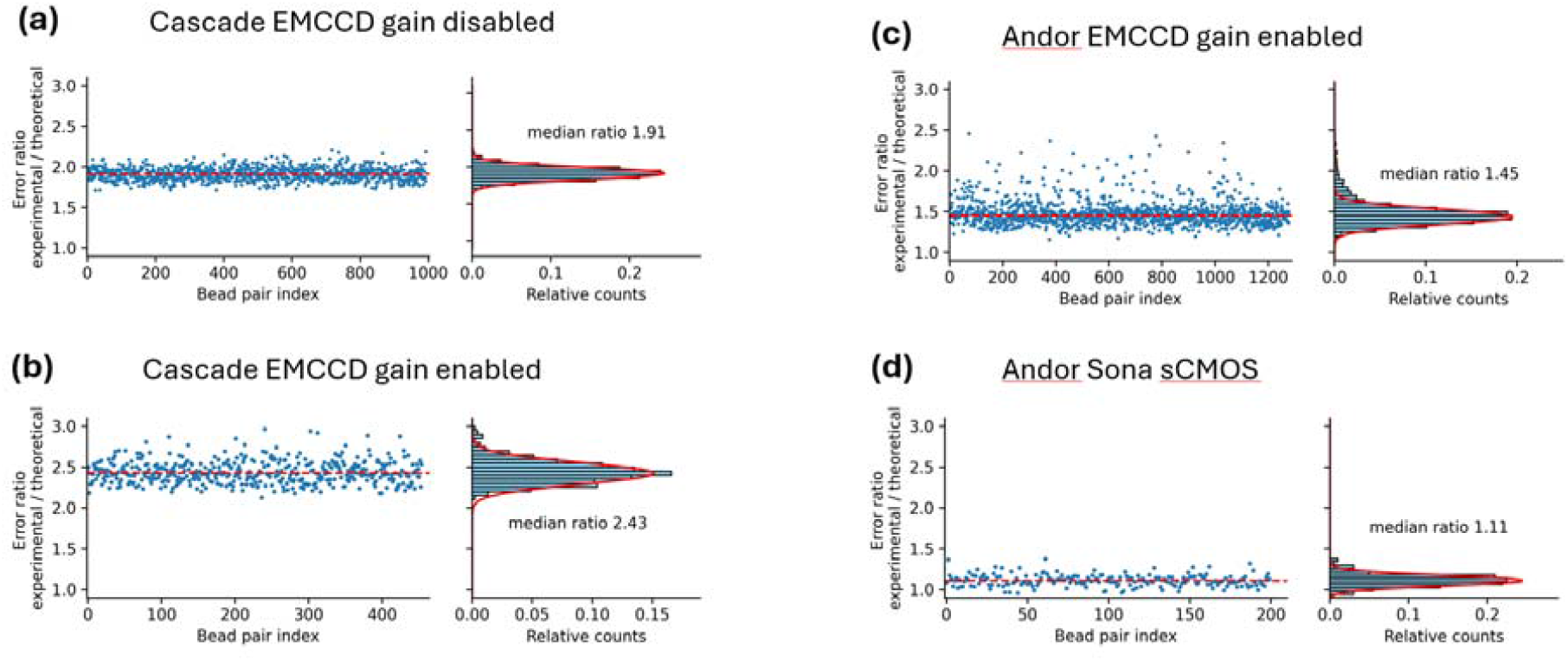
Ratio between experimental and theoretical localisation errors for different optical and detection setups. **(a)** Data for a Photometrics EMCCD camera with the gain disabled, the error ration is obtained dividing the data points in 2d. **(b), (c)** and **(d)** respectively depict the error ratios for a Cascade with the gain enabled, an Andor EMCCD and an Andor Sona sCMOS.

Interestingly, a more modern EMCCD camera (Andor iXon897) with comparable pixel size and emGain enabled (**Supplementary Table 1**), presents lower σ_exp_ / σ_theo_ ratio with a value of ∼1.5 (**Figure 3c**). Overall, these results suggest that at least part of the excess experimental error arises from the camera electronics, albeit not from the actual electron multiplication process per-se.

This appears supported by the fact that the use of an sCMOS sensor, which showed the lowest of σ_exp_ / σ_theo_ ratios, with a value of ∼1.1 (**Figure 3d**). However, it has been suggested that a different treatment of noise and calibration of the sensor individual pixels is necessary for optimal fit of sCMOS data(Huang et al. 2013). As expected, both theoretical and experimental inter-bead distance errors are inversely proportional to the total number of detected photoelectrons (**Figure S3a-b**), but the σ_exp_ / σ_theo_ ratio does not present any clear correlation with the number of photoelectrons **Figure S3c**.

The range of σ_exp_ / σ_theo_ ratios exhibited by different camera sensors presented in this work, as well as the observation that they are all larger than one, highlights the pitfalls associated to using formulas for the localization precision that strive to approximate the CRLB, as this appears to routinely lead to an underestimation of the effective localization error. Other approaches, which for example rely on the use of a PSF model calibrated on the actual setup, would allow in principle a more precise estimation of localization precision, but rely on ad-hoc software(Li et al. 2018; Liu et al. 2024), and may not consider other sources of error, such as instabilities of the setup, and in particular of the emission optics. We tested the software reported in Li et al(Li et al. 2018), which fits the single molecule peaks to the experimental PSF of the system determined from a set of calibration beads, and measured a σ_exp_ / σ_theo_ ratio of the order of ∼2.9 for the Cascade EMCCD with emGain enabled, suggesting that, also in this case, the CRLB estimate substantially underestimates the actual localization error of the system (**Figure S4**). These results overall indicate that an accessible method to provide a software-independent estimation of the localization precision of a setup, given any arbitrary code and pipeline for molecular localization, is still somehow missing.

We argue here that a straightforward procedure is a calibration for each microscope that determines the σ_exp_ / σ_theo_ ratio. To that end, we have developed a basic pipeline: in brief, a set of sub-diffraction limit fluorescent beads is imaged repeatedly over time (**Figure 1a**). Then, each bead is fit with the software of choice that the researcher desires to use in their subsequent localization microscopy experiments (**Figure 1b**). The standard deviation of the distance between each bead pair is then calculated (**Figure 1d**), and this value (σ_exp_) is compared to the expected standard deviation (σ_theo_) obtained by using error propagation from the individual beads localization precision obtained using the formula of choice (e.g. Thompson or Mortensen) (**Figure 1c**). This basic procedure which is aided by open-source analysis pipeline that relies on standard Python that furthermore does not require a GPU. Given that the experimental error is directly proportional to the experimental one (**Figure S2**), once the analysis is finalized, the computed σ_exp_ / σ_theo_ ratio allows for the calculation of the real localisation precision of each bead σ^i^_theo_ obtained from the formula of choice.

A recent review by Lelek et al. (Lelek et al. 2021) identified five widely used SMLM software suites, which add to the number of individual, custom-made algorithms used by dozens of laboratories. This number is likely to grow, considering the impact of deep-learning approaches. The proliferation of ad-hoc software for analysis, and the increase in their complexity, may have reached a point where the determination of which to choose may become challenging for a researcher. Imaging technology is also evolving and technical performance of new sensors likely to change. Finally, each individual setup has its own fingerprint in terms of optical aberrations and mechanical stability. The use of the σ_exp_/σ_theo_ ratio that we have highlighted above as a figure of merit to determine the effective localization precision of a setup appears as a software- and setup-independent approach to assess in a straightforward way the effective localization precision of single emitters, a cornerstone of the growing galaxy of super-resolution microscopy approaches.

## Materials and methods

### Sample preparation

A stock solution of TetraSpeck™ Microspheres (catalogue number T7279, ThermoFisher Scientific) with 0.1 μm diameter size and concentration of 2×10^15^ particles per mL was diluted 100 times in a 10 mM MgCl_2_ solution and sonicated 15 minutes to break particle aggregates. The bead solution was placed on a clean coverslip and incubated for 20 minutes to allow particles to sediment and attach onto the glass surface. After immobilization, the solution was gently pipetted out and replaced with pure water to prevent floating beads to interfere with subsequent microscopy measurements.

### Optical setup

A red laser beam (Cobolt, CW 638 nm) was expanded and passed through the back port of an Olympus IX73 inverted microscope. A dichroic mirror (DMLP650R) redirected the laser beam towards the sample through an Olympus 60x Apo N oil objective in either widefield epi or TIRF illumination mode (**Figure S5**). The collected fluorescence emission was magnified and focused onto a camera sensor chip. The three camera models tested in this study are an Andor sCMOS model SONA-4BV6X, an Andor iXon Ultra DU-897U-CS0-#BV and a Photometrics Cascade Serial No B07M892005. The pixel size for each camera was calculated by imaging a standard 10 μm groove reticule. All optical elements were attached to a floating optical table situated in the lowest floor of the building.

### Data collection and analysis

After bead immobilization, all movies were recorded with an acquisition time of 100 ms for a length of 1,000 frames. For Andor cameras, counts were converted to photoelectrons according to the manufacturer procedure. We used the sensitivity value reported in the System Performance Sheet which accompanied each respective camera, while the sensor background or bias was measured in total dark conditions. The sensitivity, bias and Gain response of the Cascade camera was determined in house following the manufacturer indications (**Figure S2**). For bead localisation analysis, we translated Mortensen code directly into Python and run it on our data sets. This method employs a maximum-likelihood estimation (MLE) algorithm for fitting a 2D Gaussian to the experimental PSF. Since MLE methods are slow and very sensitive to initial parameters, we additionally run a faster least-squares (LS) regression analysis and then fed the optimized parameters onto the MLE pipeline. Once the PSF centroids positions are estimated, we determined the distance over time between all possible bead pairs. The experimental localisation error is the standard deviation of the inter-bead pairs distance. The theoretical localisation error for inter-bead distance was calculated through error propagation from individual bead localisation errors computed from the Mortensen formula and averaged over all frames. The code pipeline employed in this analysis, together with detailed instructions on running the software is available at *https://github.com/brenlla1/Localisation-precision*.

## Author contributions

AB-L. and P.A. designed research; AB-L. and L.D. performed research; P.A. and A.B-L. wrote the paper with input from L.D. P.A supervised the project and acquired funding.

## Acknowledgments

PA gratefully acknowledges support from the Leverhulme Trust (RL-2022-015).

## Supplementary Tables and Figures

**Supplementary Table 1.** Optical and electronic acquisition parameters for all measurements shown in this work.

| Camera model | Pixel size<br>/ nm | Pixel area | Electron<br>multiplication | Median ratio<br>$\sigma_{\text{exp}} / \sigma_{\text{theo}}$ |
| --- | --- | --- | --- | --- |
| Photometrics Cascade<br>Serial No **B07M892005** | 89 | 512 x 512 | Yes | 1.91 |
|  |  |  | No | 2.43 |
| Andor iXon Ultra **DU-897U-CS0-#BV** | 85.4 | 512 x 512 | Yes | 1.45 |
| Andor sCMOS **SONA-4BV6X** | 34.2 | 1024 x 1024 | No | 1.11 |

**Supplementary Figure 1.**
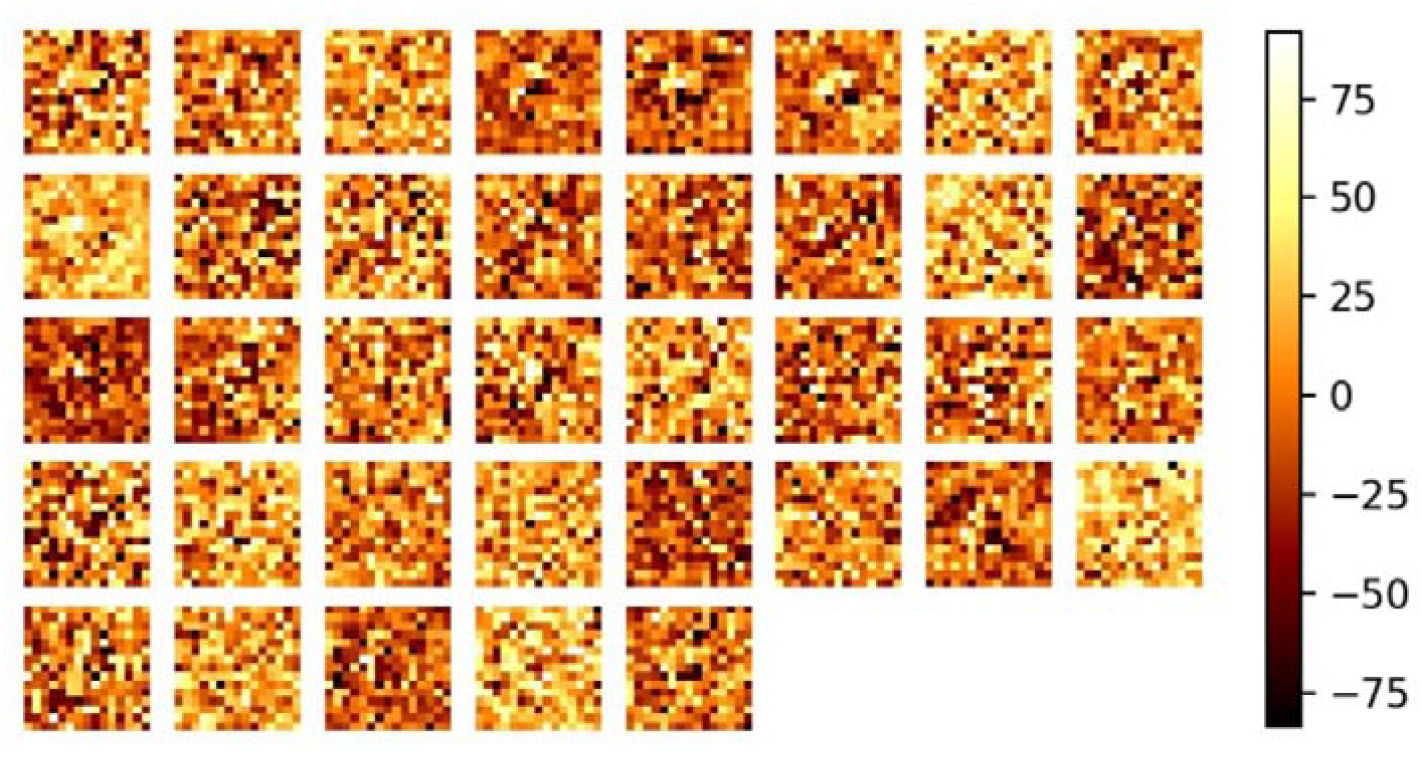
Representative residual plots from multiple beads imaged using the Photometrics Cascade camera without emGain.

**Supplementary Figure 2.**
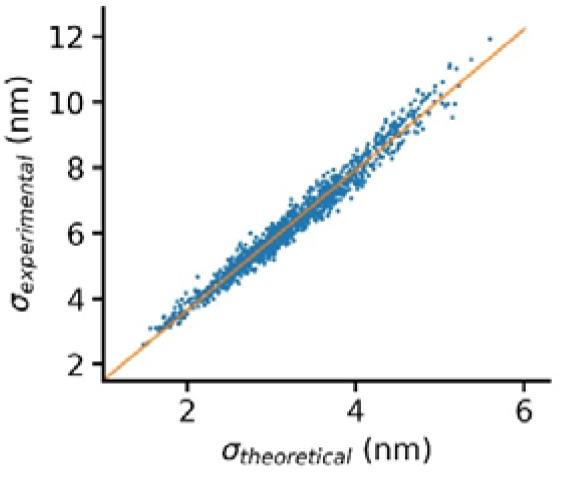
The experimental localisation error (y-axis) is proportional to the theoretical error (x-axis), the orange line corresponds to a linear fit. The data shown here were recorded in the Photometrics Cascade camera with EM Gain disabled.

**Supplementary Figure 3.**
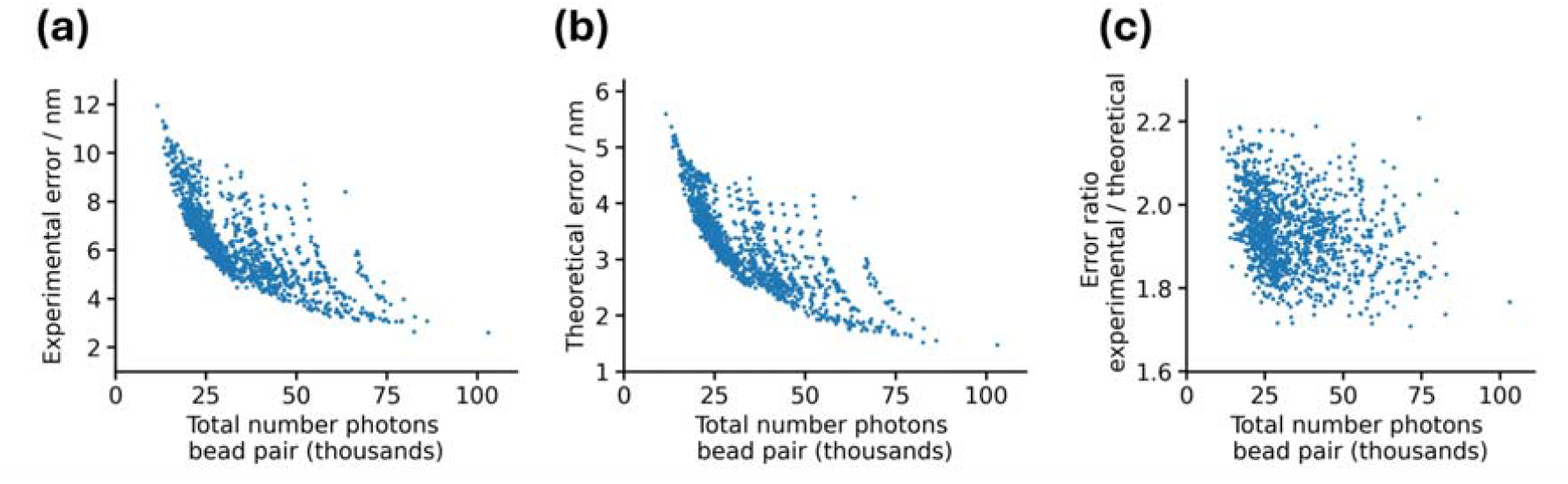
Experimental **(a)** and theoretical **(b)** errors for inter-bead separation as a function of the number of photons for the data recorded in the Photometrics Cascade with EM Gain disabled. **(c)** Ratio between the experimental and theoretical error shown in (a) and (b) respectively. The total number of photons corresponds to the sum of photons detected from each bead in each bead pair.

**Supplementary Figure 4.**
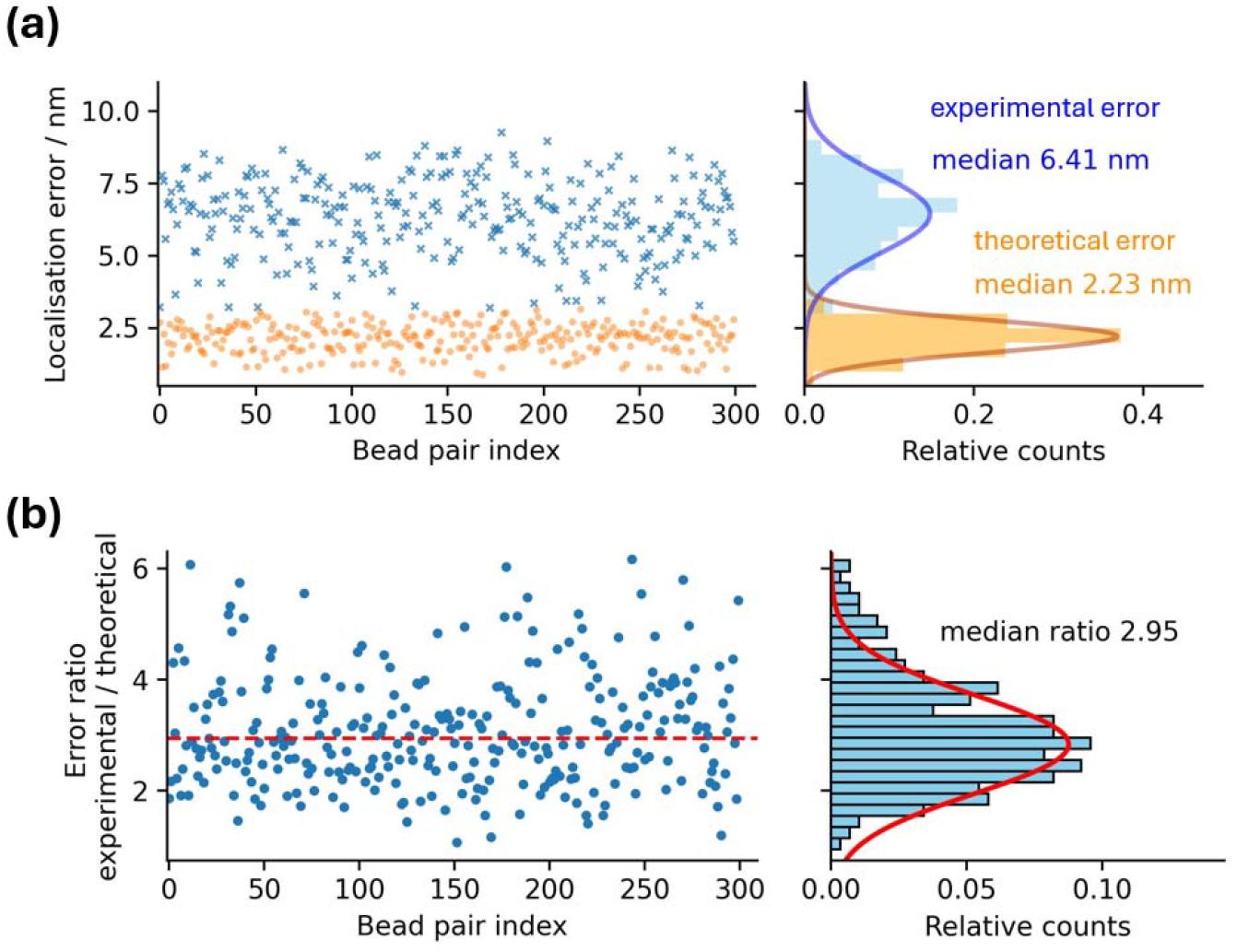
Experimental and theoretical localization errors using an experimental PSFmodel. **(a)** Experimental and theoretical localisation errors for over 300 bead pairs measured using the Cascade camera with an emGain value of 3,500.

**Supplementary Figure 5.**
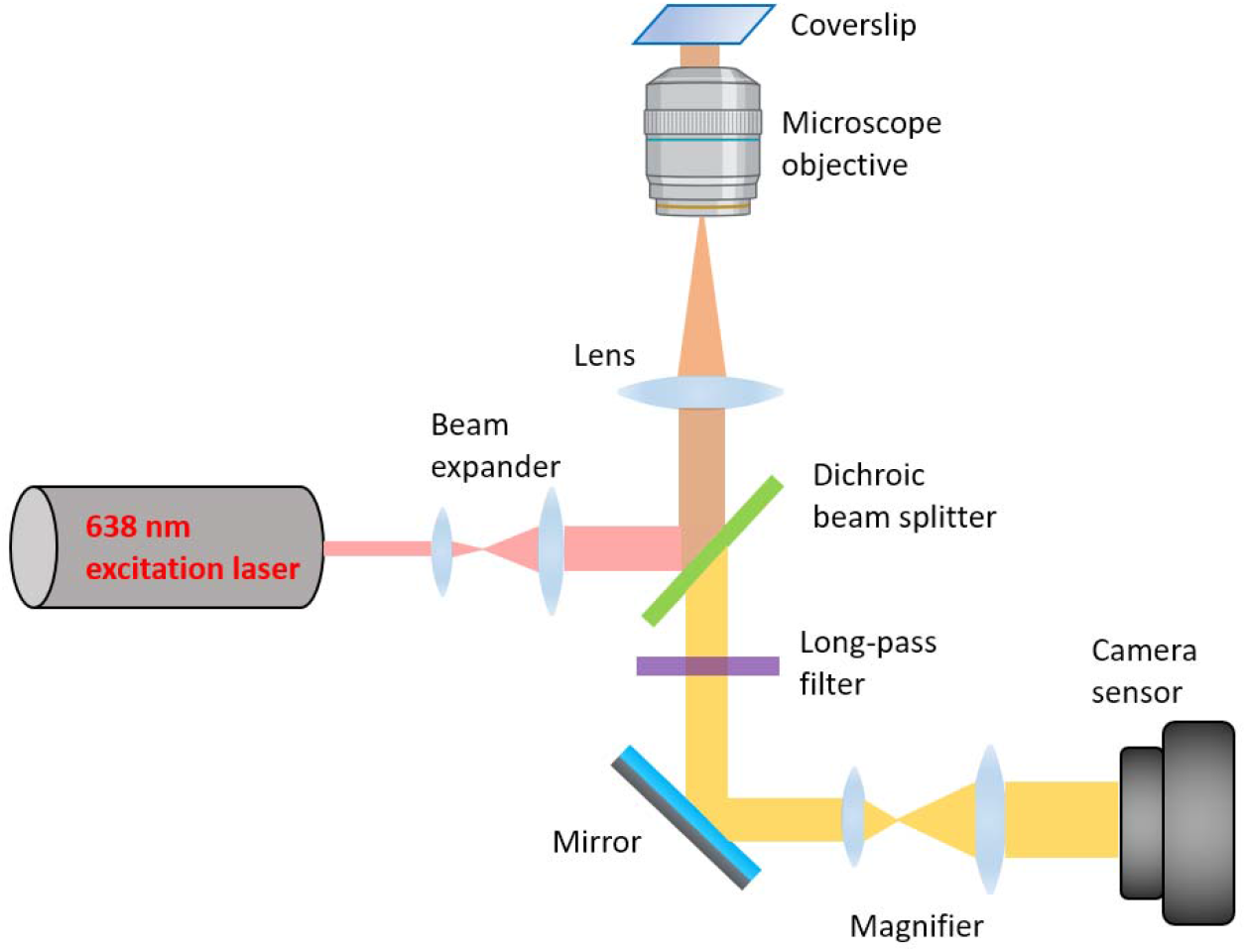
Schematic of the optical setup used in this work.

**Supplementary Figure 6.**
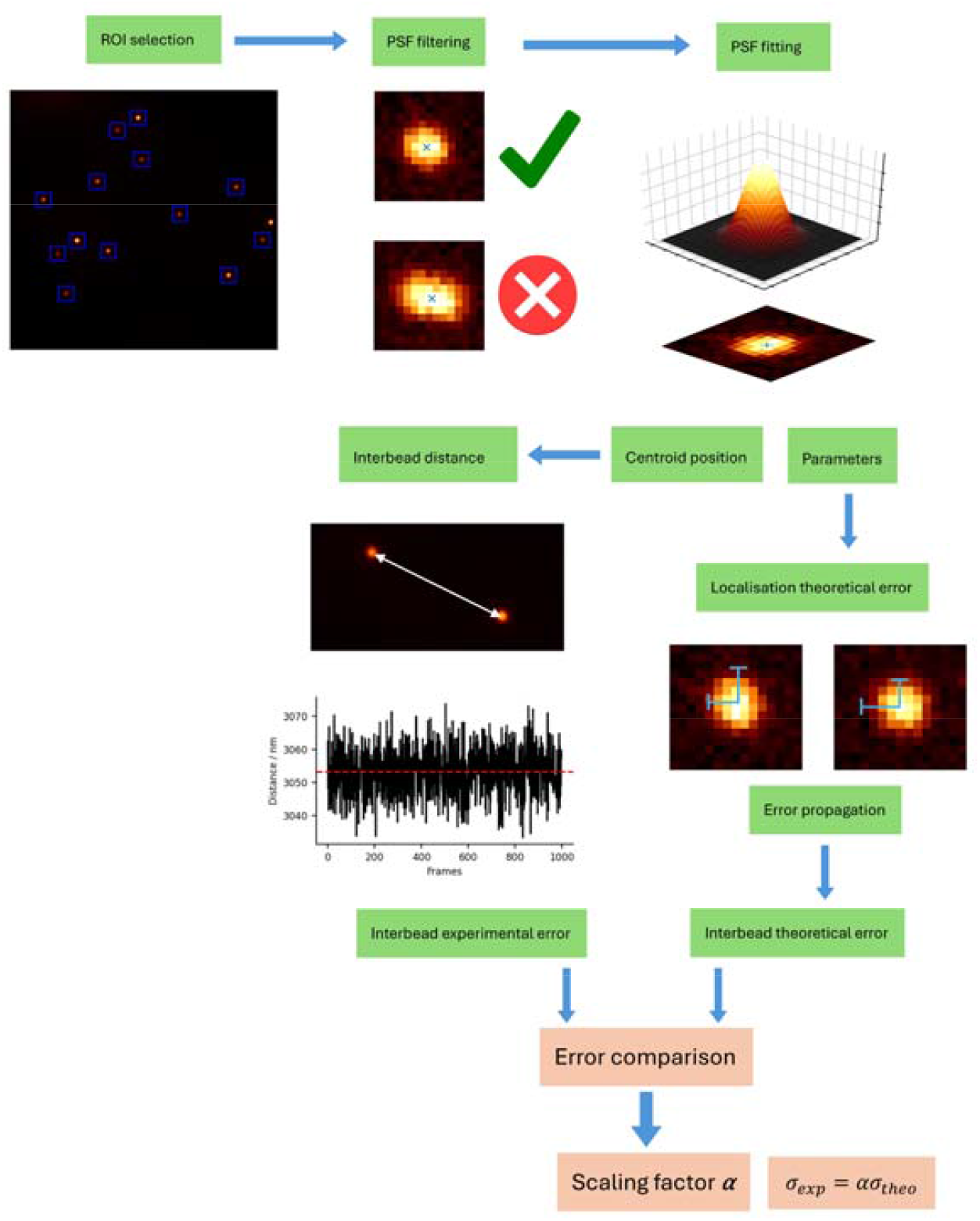
Data pipeline for obtaining both the experimental and theoretical errors.

